# Systemic antibody repertoires target specific microbial antigens in patients with colorectal cancer

**DOI:** 10.64898/2026.10.15.759395

**Authors:** Gabriel Innocenti, Stefanie Brezina, Iros Barozzi, Melanie Prinzensteiner, Carlos S. Reyna-Blanco, Mateusz F. Kolek, Julian Prosenz, Andreas Maieron, Andrea Gsur, Thomas Vogl

## Abstract

Colorectal cancer (CRC) represents the third leading cause of cancer related deaths worldwide and is influenced by multiple risk factors. Both exposure to microbiota as well as adaptive immune responses have been implicated in CRC development and suggested as candidate biomarkers. However, the extent to which CRC and adenomas associate with systemic immune signatures against microbes remains unclear.

Here, we apply Phage-display ImmunoPrecipitation Sequencing (PhIP-Seq) on 321 individuals of an Austrian CORSA cohort (Colorectal Cancer Study of Austria) including controls, adenomas, and CRC patients. Although antibody repertoires showed global similarity across the different disease status and limited predictive performance, prevalence-based analyses revealed specific signals, such as *Helicobacter pylori* CagY as candidate marker of CRC and viral antigens as candidate for non-cancer stages. Taken together, this large-scale analysis of antibody reactivity in CRC highlights the potential of serological markers to reflect disease-associated immune signatures.

## Introduction

Colorectal cancer (CRC) is the third leading cancer related cause of death worldwide and represents a major public health issue (1). It is characterized by several established modifiable and unmodifiable risk factors such as age, male sex, family history of CRC, genetic factors, inflammatory bowel disease, diabetes, obesity, smoking, physical inactivity, alcohol consumption and high consumption of red meat (2). Most CRCs are sporadic and usually develop in a slow progression from normal epithelium to precancerous adenomas to invasive cancer. This trajectory offers significant opportunities for preventive screening and early intervention through the effective detection and removal of precancerous adenomas, thereby reducing mortality (3).

In the recent years, studies have shown that dysbiosis of the gut microbiota is linked to CRC development and progression (4). In particular, proliferation of specific genotoxin-producing bacteria such as pks+ *Escherichia coli* (5), as well as pro-tumorigenic taxa as *Fusobacterium nucleatum* (6), and *Peptostreptococcus anaerobius* (7) were associated with an increased risk of developing cancer. Several studies have reported signatures in microbiota associated with an increased risk of both CRC and pre-cancerous stages (8). Most of these efforts have relied on Next Generation Sequencing technologies of fecal samples, including 16S amplicon sequencing and shotgun metagenomics, to large study cohorts (9–12). Together with the growing interest in machine learning approaches, these efforts have also led to the development of microbiota-based predictive models for CRC prediction and risk stratification (13,14).

It has generally been appreciated that not the presence of specific microbes, but rather their interaction with the human immune system could influence pre-cancerous and cancerous stages (15). As part of this microbiota-immune axis, microbes modulate the immune system through exposure of immunoreactive antigens and production of toxins and metabolites. Specifically, these microbial antigens can elicit both local (mucosal) and systemic adaptive immune responses, including circulating antibodies that recognize components of microbiota (16,17). Therefore, insights into cancer etiology and new diagnostic approaches may derive not only from the presence and abundance of certain strains in the gut microbiota, but also from the immune responses they elicit systemically and locally. However, given that there are thousands of bacterial strains with thousands of genes encoding potential antigens (18), immune responses against the microbiota are difficult to map with conventional methods such as ELISAs or peptide arrays (19).

Phage-display ImmunoPrecipitation Sequencing (PhIP-Seq) allows characterizing antibody responses against hundreds of thousands of antigens in hundreds to thousands of individuals (20–22). This method served multiple purposes, including studying antibody repertoires against microbial structures in ∼1,000 healthy individuals (23), inflammatory bowel disease (IBD) (24), and primary sclerosis cholangitis (PSC) (25). However, whether CRC and adenomas are associated with distinct antibody repertoires against microbiota-derived antigens has never been tested on a large scale.

Here, we applied PhIP-Seq to detect systemic immune responses of 321 individuals recruited within the ongoing “Colorectal Cancer Study of Austria” (CORSA) (26), including colonoscopy-negative controls, adenomas and patients diagnosed with CRC. We tested samples against an antigenic library composed by ∼357,000 antigens, composed of three previously published PhIP-Seq libraries (23,27,28).

## Results

### Antibody repertoires are globally shared across controls, adenoma and CRC

We performed PhIP-Seq on 321 plasma samples from patients recruited within CORSA (see Methods), screening each patient for 357,192 antigens included in three previously published PhIP-Seq libraries (Fig. 1A, Methods). Our study cohort was divided into three outcome groups: Controls (N=107), Adenoma (N=107) and CRC (N=107), which were matched for age and sex to rule out potential confounding effects. After sequencing, antibody reactivity profiles for each patient were obtained by calculating the number of enriched antigens as outlined in the Methods section. We removed 10 samples which showed no enriched antigens (having likely failed due to experimental issues) yielding a final pool of 311 samples (N_Controls_= 102, N_Adenoma_ = 105, N_CRC_= 104), subsequently used for statistical analysis (Supplementary Table 1). Age/sex matching was not affected by removing these samples (Fig. 1B). Also, because the exact time point of sample recruitment could be a confounding factor for massively parallel serology, we verified that the year of recruitment was evenly distributed across the clinical groups (Supplementary Figure 1A).

**Fig. 1:**
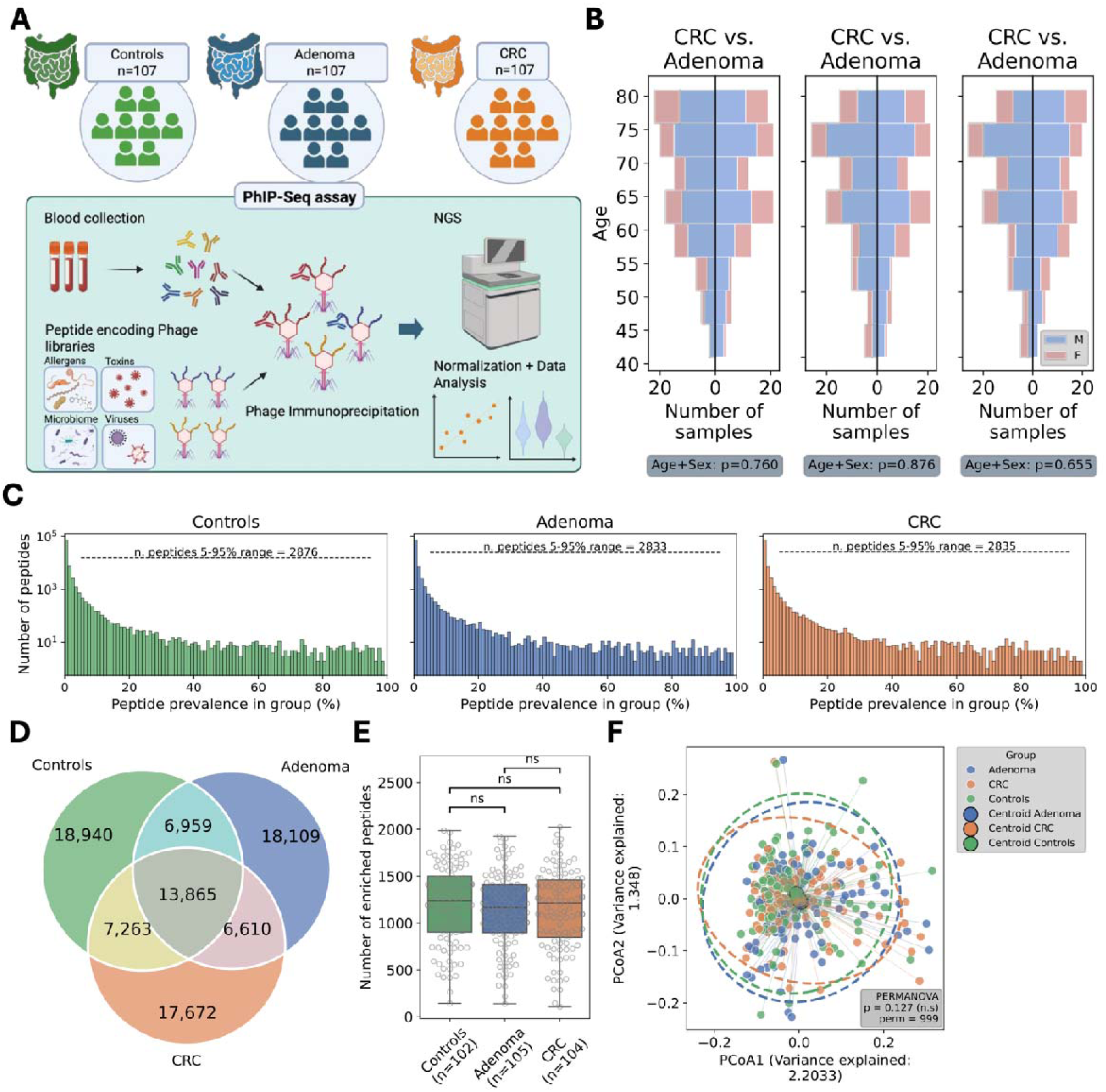
Phage immunoprecipitation sequencing shows that antibody repertoires maintain global compositional stability across three disease groups consisting of CRC, adenoma, controls. **A.** PhIP-Seq workflow allows to screen antibodies against ∼357,000 different antigens from several sources including human microbiota, allergens, human viruses, pathogenic strains and autoantibodies. **B.** Age and sex distributions are consistently matched across all the groups in this study. **C.** Reactive antigens distribution profiles showing the number of peptides that are bound for each prevalence interval in each group (Controls, Adenoma, CRC). **D.** Venn plot showing the antibody bound antigens which overlap across the three groups of patients and the unique antibody repertoires specific to each group. **E.** Number of enriched antigens distribution across each group, compared using Mann-Whitney test. **F.** Principal coordinate analysis using antigens presence/absence data on the full dataset and colored by groups. Centroids were calculated as mean of distances within each group and ellipses were added represent ∼95% concentration of data points for each group.

Continuing with 311 samples, we detected in total n = 89,418 enriched antigens. We observed individual specific responses in all three groups, as well as shared immune responses across high percentages of patients within each group (Fig. 1C). We observed a core peptidome for the three groups consisting of 13,865 antigens present in at least one sample for each group, while a higher number of antigens were disease specific (Controls = 18,940; Adenoma = 18,109; CRC = 17,672, Fig. 1D). Given that the antibody reactivity is largely dominated by individual-specific antigens or shared among low fraction of individuals per group, we restricted the analysis to antigens shared within the 5-100% prevalence range in each group. In this case, the core peptidome included most antigens, while the group-specific antigens proportion decreased (Supplementary Figure 1B), indicating a large, shared antigen pool between the different disease groups. This observation of private and public repertoires is also in line with previous findings in healthy individuals (23,29) and disease (24).

We also observed overall similar numbers of antigens bound per sample in each group, showing no significant difference (Fig. 1E). In the same way, alpha diversity, as a proxy of intra-sample antigen diversity, showed consistent stability across each group (Supplementary Figure 1C). We performed dimensionality reduction and Principal Coordinate Analysis (PCoA) on the whole enriched antigens pool. Variance explained by each dimension was relatively low (including the first 2: Var_PCoA1_=2.203, Var _PCoA2_=1.348), and PERMANOVA test detected no significant difference among groups (pvalue=0.131, permutations=999) (Fig. 1F, Supplementary Figure 1D). As expected, age was a significant contributor to the explained variance (PERMANOVA pvalue=0.001), while sex showed a modest but also significant contribution (PERMANOVA pvalue = 0.045). Similar results were also found after aggregating the antigens data to organism species, which also showed no significant difference between groups (Supplementary Figure 1E). To explore the relationship between the data and clinical outcomes, we tested all possible prevalence combinations using a lower threshold ranging from 0 to 30% and an upper threshold from 70 to 100%. For each filtered antigen set, we performed PERMANOVA test to assess clinical group associated differences. The resulting p-value map showed that reducing antigenic space to given prevalence intervals increased the association between antigens presence/absence and the clinical outcome (Supplementary Figure 1F). However, even with the most significant antigen subsets we could not distinguish clear clusters when comparing the first PCoA components, indicating that the variance explained remained evenly distributed across all dimensions (Supplementary Figure 1G). These results illustrate that antibody repertoires showed high-level similarity between controls, adenoma and CRC.

### Pairwise comparisons across groups reveal differentially prevalent antibody responses

High level metrics (Fig. 1) showed no clear difference between CRC, adenomas, and controls, so we investigated differences for specific antigens and species. Most of the enriched antigens lay in a very low prevalence interval (Fig. 1C) and therefore did not carry group-wise but rather individual-specific information. For this reason, we selected antigens present in at least the 5% of each clinical group and compared antigenic prevalences pairwise between groups. Following a Fisher test we observed specific antigens which were significantly more prevalent either in one or the other clinical group. However, no antigens remained statistically significant after False Discovery Rate (FDR) correction, likely due to the high number of comparisons performed. We evidenced the different peptides percentage distribution within each comparison in the scatterplots in Fig. 2. When comparing CRC vs. controls (Fig. 2A), the top hits for CRC were represented by viral proteins (*Cytomegalovirus* and *Simplexvirus humanalpha 4*) and most notably several hits of the Cytotoxic immunodominant antigen belonging to *Helicobacter pylori* (Fig. 2B, Supplementary Table 2). For controls, most differentially prevalent antigens belonged to EBNA1 (Epstein-Barr virus), Hemagglutinin (*Betainfuenzavirus influenzae*), endo-beta-N-acetyl glucosaminidase (*Streptococcus pneumoniae)* and viral polyproteins (Norwalk virus and Rhinovirus A) (Fig. 2B, full data in Supplementary Table 2). This underlines a higher presence of antigens belonging to human viruses in controls, especially viruses such as *Betainfluenzavirus influenzae* and *Rhinovirus A* which are commonly causative of seasonal flu.

**Fig. 2:**
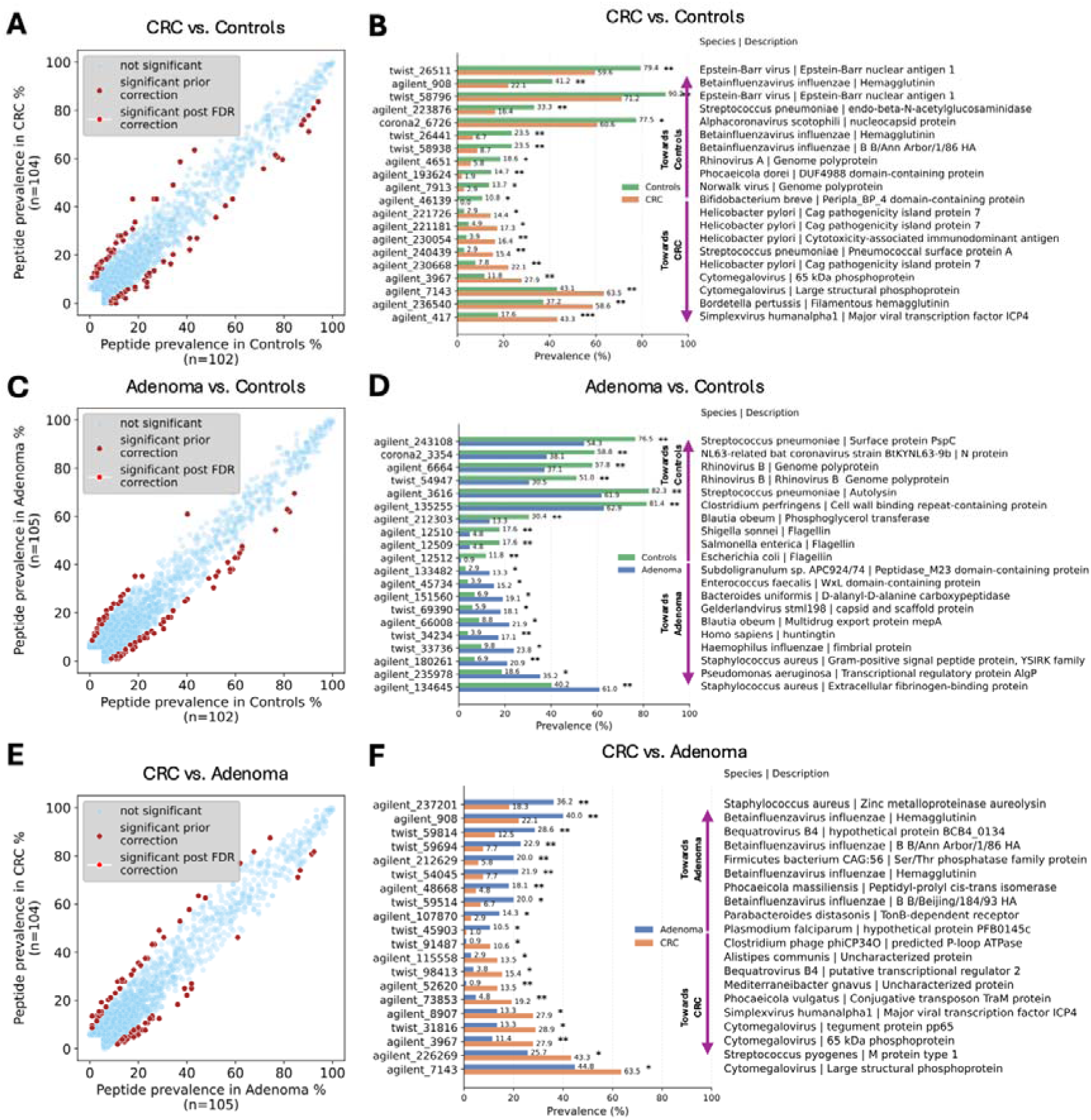
Comparing antibody responses in 5-100% of each group shows no mayor prevalence shifts but shows significant hits after running Fisher’s test on the prevalence of each peptide across each group comparison. **A.** Scatterplot showing for each peptide (single dots) the ones that are differentially prevalent between the groups of controls and CRC using Fisher’s exact test to assess significance. **B.** Bar plot showing the top reactive antigen hits differentially prevalent in CRC or controls (* = 0.05, ** = 0.01). Description of antigens was added including species and protein name. **C.** Scatterplot showing differentially prevalent peptides within the comparison controls vs. adenoma. **D.** Bar plot showing the top reactive antigen hits differentially prevalent in controls or adenoma. **E.** Scatterplot showing differentially prevalent peptides within the comparison adenoma vs. CRC. **F.** Bar plot showing the top reactive antigen hits differentially prevalent in adenoma or CRC.

Comparing adenoma versus controls also revealed different trends (Fig. 2C). Adenoma showed a higher presence of extracellular fibrinogen-binding protein and signal peptide (*Streptococcus pneumonia*e) but also higher antigenic proteins coming from pathogenic strains such as *Pseudomonas aeruginosa*, *Enterococcus faecalis*, *Listeria monocitogenes* and *Haemophilus influenzae* (Fig. 2D). In contrast, controls were enriched in *Streptococcus pneumoniae* and *Rhinovirus B* polyproteins, but also flagellins and cell-membrane proteins derived from taxa commonly associated with the human microbiota, including *Escherichia coli*, *Shigella sonnei*, *Clostridium perfringens* and *Salmonella enterica* (Fig. 2D).

Finally, we observed some differences between CRC and adenoma antibody repertoires (Fig. 2E). In cancer we observed the most prevalent hits highlighting Cytomegalovirus structural proteins, as well as other viral proteins belonging to Simplex virus and Varicella virus. Other relevant proteins came from microbiota strains such as *Streptococcus pyogenes*, *Alistipes communis* and *Phocaeicola vulgatus* (previously known as *Bacteroides vulgatus*) – both belonging to Bacteroidales order (Fig. 2F). The most prevalent hits in adenomas were mostly represented by Betainfluenzavirus influenzae Hemagglutinin protein (from different strains), while also showing other more prevalent antibody responses against *Firmicutes bacterium CAG:56*, *Phocaeicola massiliensis* and *Parabacteroides diastonis* compared to CRC (Fig. 2F). Interestingly, this did not emerge in the previous comparison.

Overall, these results point to a shift in antibody repertoire patterns across the three groups. Interestingly, among the most prominent signals we could highlight *Betainfluenzavirus influenzae* Hemagglutinin consistently as relevant in Adenoma against Cancer and, on the other side, immunogenic antigens such as CagA were more prevalent in CRC when compared to controls.

### Species and protein-level aggregation show disease-associated antibody responses driven by immunodominant antigens

While none of the single peptides observed were significantly different between groups after correction for multiple hypothesis testing (Fig. 2), several peptides originated from the same species. This finding suggests, that while exact epitopes have limited power, immune responses against species may carry additional information for biomarker discovery. Hence, we applied a statistical framework to aggregate antigen prevalence to the corresponding species level, testing for significance in each clinical group pair comparison (Methods). When comparing CRC versus controls, we found a strong signal of antibodies targeting *Helicobacter pylori* (n. peptides = 271, p = 0.006) showing higher prevalence in CRC, while in controls we found *Betainfluenzavirus influenzae* as the main hit (n. peptides = 124, p = 0.0002). Other strains such as *Bifidobacterium breve* (n. peptides = 287, p = n.s.) and SARS-CoV (n. peptides = 344, p=n.s.) were characterized by strong effect size but did not show significance (Fig. 3A, Supplementary Table 3).

**Fig. 3:**
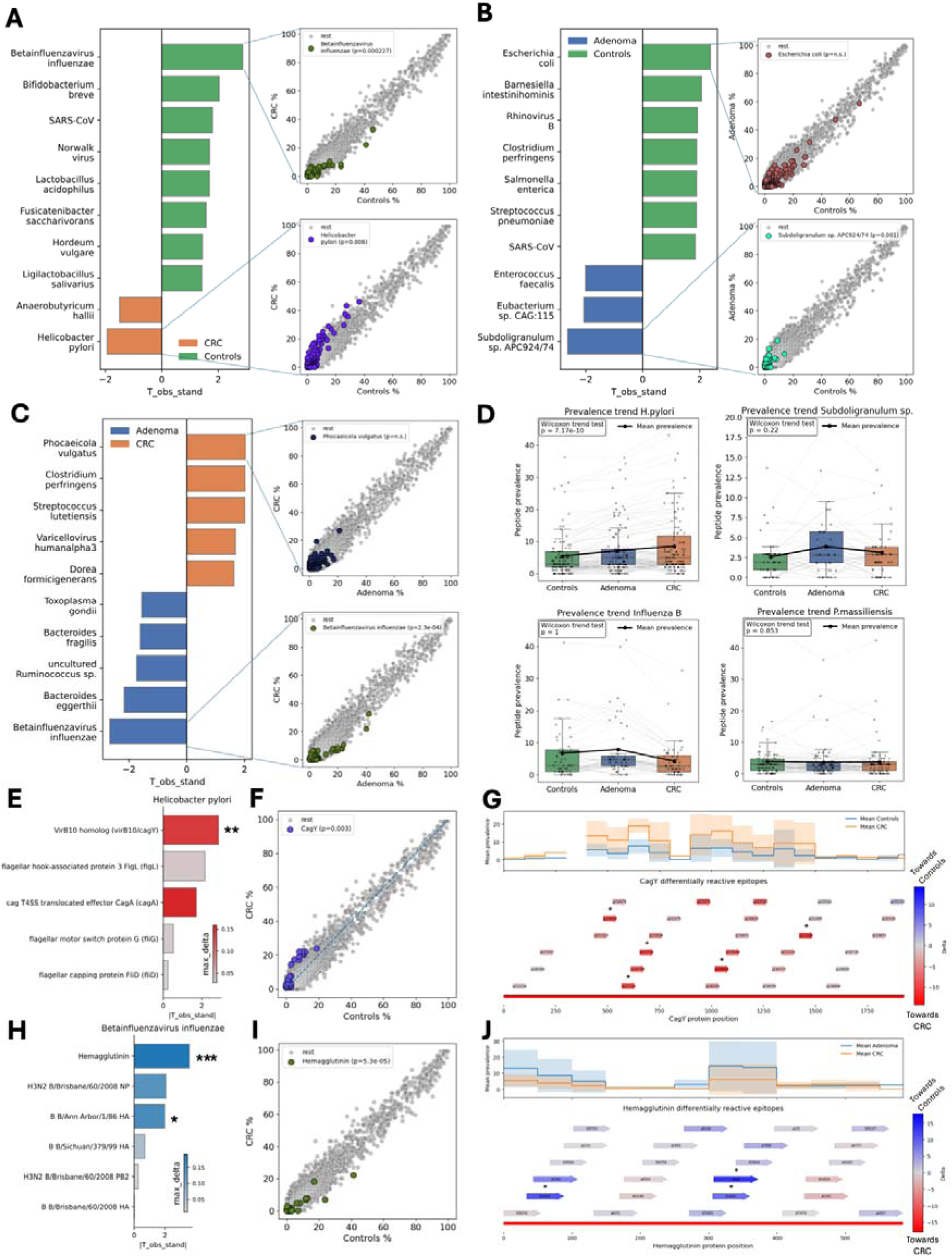
Antigens aggregated prevalence shows the presence of differentially prevalent species and protein across the three group comparisons. **A.** Bar plot showing the top ranked species in aggregated prevalence in CRC vs. controls in each direction. We applied a filtering according to effect size (>30 used peptides), T standardized coefficient (>1.5) and maximum prevalence delta (>0.1) between the two groups. Prevalence distribution for the best hit in each direction is shown in the prevalence scatterplots, coloring the antigens mapped on the given species. **B.** Bar plot showing the top ranked species in aggregated prevalence in adenoma vs. CRC. Prevalence scatterplots for the best hit in each direction also shown. **C.** Bar plot showing the top ranked species in aggregated prevalence in adenoma vs. controls as explained in A. Prevalence scatterplots for the best hit in each direction also shown. **D.** Mean reactivity score was calculated averaging the protein reactivity score (Methods) within each species per sample and was compared across groups. Both a Spearman correlation test and linear regression were used to assess whether an increasing or decreasing trend from Controls to Adenoma to CRC was present. **E.** *Helicobacter pylori* top ranked proteins according to T standardized statistic when peptide aggregation was done at protein level. **F.** Prevalence scatterplot showing *Helicobacter pylori* CagA-derived antigens prevalence distribution compared to the rest. **G.** Mean prevalence difference showed along all the peptide tiles covering CagA protein. **H.** *Betainfluenzavirus influenzae* top ranked proteins according to T standardized statistic when peptide aggregation was done at protein level (* pvalue < 0.05; ** pvalue < 0.01; *** pvalue < 0.001). **I.** Prevalence scatterplot showing *B. influenzae* hemagglutinin-derived antigens prevalence distribution compared to the rest. **J.** Mean prevalence difference shown along all the peptide tiles covering hemagglutinin protein from *B. influenzae*.

When comparing CRC to adenoma we found *Betainfluenzavirus influenzae* as a main hit as well (n. peptides = 128, p=2.4x10^-4^), along with other non-significant strains, that showed an increase in prevalence within the adenoma group (Fig. 3B, Supplementary Table 3). No significant strains were found in CRC, although *Phocaeicola vulgatus* (n. peptides = 2,153, p= n.s.) was slightly more prevalent in CRC compared to adenoma. Lastly, we compared adenoma and controls, observing a higher prevalence of some species in adenoma, including *Escherichia coli* (n. peptides = 1,785, p= n.s.), *Clostriudium perfringens* (n. peptides = 526, p= n.s.) and *Salmonella enterica* (n. peptides = 917, p= n.s.), characterized by a relevant effect size and T coefficient although not showing any significance.

On the other hand, controls showed an increased prevalence of Subdoligranulum sp. APC/924/74 (n. peptides = 262, p= 0.001) and non-significant hits for Eubacterium sp. CAG:115 (n. peptides = 44, p = n.s.) and Enterococcus faecalis (n. peptides = 464, p = n.s.) (Fig. 3C, Supplementary Table 3).

These results highlight *Betainfluenzavirus influenzae* as a potential immune system target characterizing non-cancerous stages, while *Helicobacter pylori* as possible immunogenic indicator of CRC. A more varied signal for controls consisting of higher presence of microbiota strains belonging to *Subdoligranolum*, *Eubacterium* and *Bifidobacterium* genera.

We took the main taxonomic hits identified in each comparison and evaluated whether we could identify a progressive increasing or decreasing trend in the detected antigens prevalence from controls to adenoma to CRC. Helicobacter pylori was the only candidate which showed a clear increasing trend (Wilcoxon trend test p = 7.17*10^-7^), while for *Phocaeicola vulgatus*, *Betainfluenzavirus influenzae* and *Subdoligranulum APC924/74* no visible trends were observed (Fig. 3D). Furthermore, we computed a mean antigenic reactivity breadth on these same strains, to test whether the antibody responses were driven by a small or high proportion of antigens for each species and we repeated the trend test (see Methods, Supplementary Figure 2). Also in this case, only *Helicobacter pylori* showed an increasing trend (Spearman p = 2.23 x 10^-2^; Linear regression p = 2.08 x 10^-3^, R^2^=0.04), from controls to CRC, while generally stable trend was found for in *Phocaeicola vulgatus*, *Betainfluenzavirus influenzae* and *Subdoligranulum APC924/74*.

Since we found significant candidates in both directions in separating CRC and controls, we focused on this comparison and specifically on *Helicobacter pylori* as a representative species for CRC and *Betainfluenzavirus influenzae* as representative for controls. We then repeated the statistical aggregation analysis at the protein level and ranked the results based on the standardized T coefficient (Fig. 3E). Interestingly, CagY and CagA, proteins belonging to the Cag pathogenicity island, were among the most prominent hits for CRC (Supplementary Figure 3A-E). CagY showed the highest aggregated prevalence difference among *Helicobacter pylori* antigens, and the only significant hit (n. peptides = 52, p=0.003; Fig. 3F). CagY entire protein is 1,927aa amino acid-long and we found antibody responses against the whole protein sequence. Comparing the prevalence profile of each reactive antigen mapping on CagY, we could recognize two regions of the protein - pos_1_: 300-700aa and pos_2_: 900-1500aa - where the difference in prevalence between CRC and controls was maximized (Fig. 3G). This suggests that the antibody response in CRC and controls is directed toward largely overlapping regions of CagY, but with a higher prevalence of reactivity among CRC patients.

Similar behavior was observed for Hemagglutinin protein from *Betainfluenzavirus influenzae*, which was the most prominent hit based on the T standardized statistic (n. peptides = 31, p=5.3 x 10^-5^; Fig. 3H,I). Hemagglutinin is a 585-aa-long protein present in most of Influenza viruses: Here, the region spanning between 300-400aa was subjected to higher antibody reactivity in controls compared to cancer patients, while reactivity across the rest of the protein remained broadly similar in the two groups (Fig. 3J). We observed that antigens being more prevalent in controls were mapping on a conserved region of Influenza B virus strains, probably due to a previous exposure to the virus or vaccination (Supplementary Figure 3F,G).

Although not showing a strong pattern difference, these results suggest the presence of potential species-specific and protein-specific candidates for each group, especially in CRC and controls.

### Antibody profiles show slight predictive capacity detecting cancer against non-cancer stages

Given the large number of enriched antigens and balanced sample size across the groups, we performed machine learning to evaluate the predictive potential of the enriched antigens as biomarkers. Specifically, we ran a random forest classification model (explained in the Methods) to assess whether it was possible to predict the disease status based on the antigen reactivity presence-absence profile in our dataset. We used a nested cross-validation setup and inner-loop feature selection to remove the noise from our dataset and a random forest classifier to perform the predictions. The random forest models were trained for all group pairwise comparisons: CRC versus controls, adenoma versus controls and CRC versus adenoma (Fig. 4A). We additionally tested predictive performance aggregating controls and adenoma together as “non-disease” versus CRC, and adenoma and CRC versus controls.

**Fig. 4:**
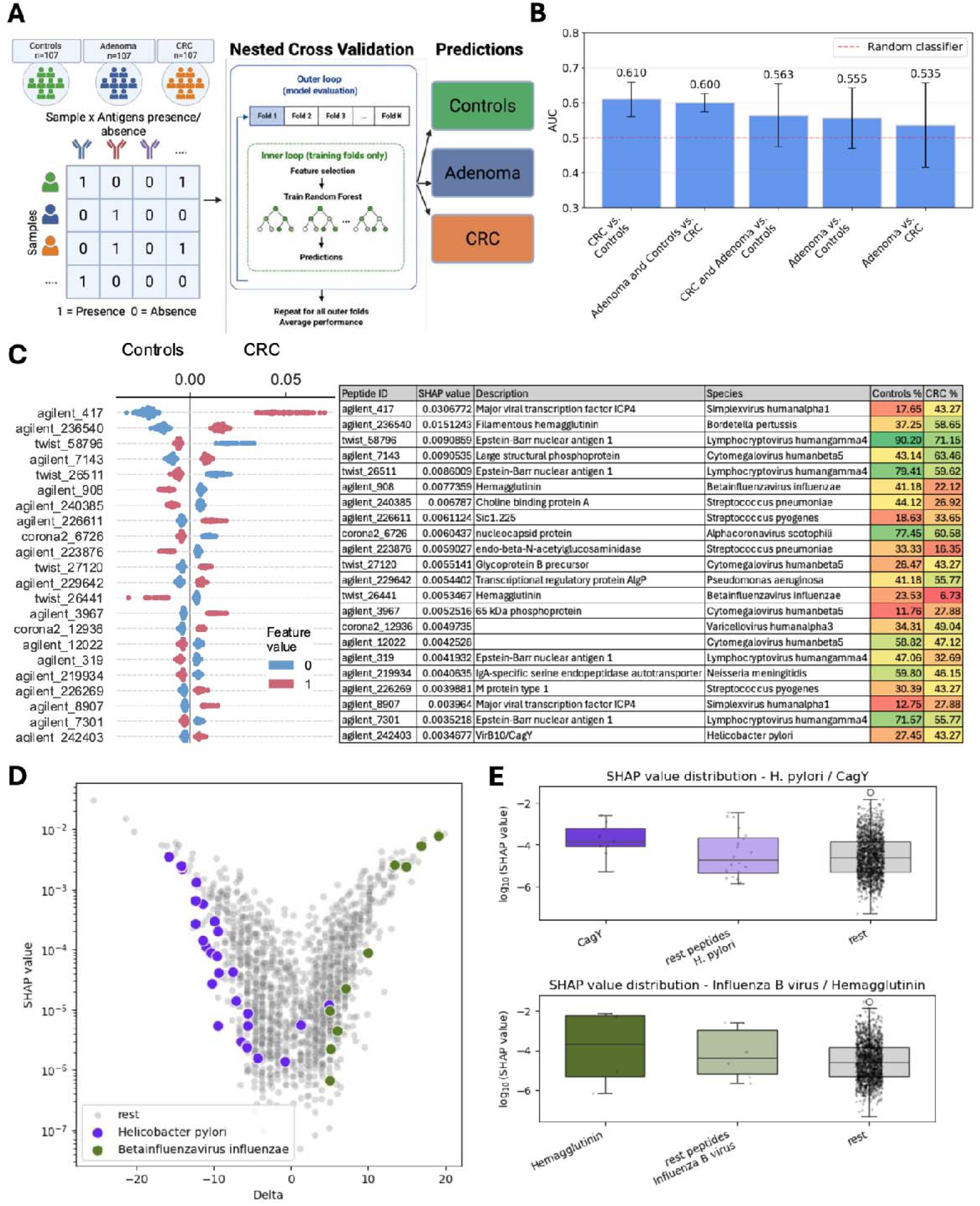
Predictions using presence-absence of antigen reactivity show moderate performance in accurately classifying the disease groups, with *Helicobacter pylori* and *Betainfluenzavirus influenzae* antigens consistently detected as top features. **A.** Classification model schema consisting of using the antigens presence/absence matrix from the three groups as input for a nested cross-validation random forest classifier, with model selection and hyperparameters tuning performed within the inner loops. Outer loops are used to evaluate the model and give a more confident prediction avoiding data leakage. **B.** Comparison of AUC performance for the model classification based on pairwise groups comparisons; The 95% confidence interval levels are also shown in the plot for each model. **C.** SHAP feature contribution plot for the CRC versus controls model; It shows both the direction and importance of each peptide within the model. On the right, a table showing the information relative to the first 22 features (antigens) appearing in the SHAP plot. **D.** Relationships between SHAP value and antigens delta prevalence between CRC and controls, mapping the antigens belonging to H. pylori and *Betainfluenzavirus influenzae*. **E.** Boxplot showing different distributions of SHAP values (in logarithmic scale) between the CagY protein from *H. pylori*, the other antigens belonging to this Bacteria and all the rest of the antigens (we excluded here the SHAP values equal to zero). F. Same boxplot showing the distribution of SHAP values comparisons between Hemagglutinin, other antigens derived from *Betainfluenzavirus influenzae* and all the remaining antigens.

Overall, we observed a modest performance across all the classification tasks. The highest performance was found when predicting CRC versus controls, with an AUC = 0.61 (95% CI = [0.561-0.659], ±1SD=0.04), followed by CRC versus both adenoma and controls, showing an AUC=0.60 (95% CI = [0.574-0.626]) ±1SD= 0.021) (Fig. 4B, Supplementary Figure 3A). The remaining comparisons showed a lower performance close to random classification, indicating a limited discriminative power (Supplementary figure 3B-E). We determined the contribution of each antigen feature by calculating SHAP values (Fig. 4C, Supplementary Figure 3A-E). Considering the best performative model (CRC vs. Controls) we could observe that the mainly contributing antigens were overlapping with the ones found to be differently prevalent in the given comparison (Fig. 2A-B), highlighting the importance of Cytomegalovirus, Simplex virus alpha and Streptococcus genus (Fig. 4C). We extracted the features belonging to *Helicobacter pylori* and *Betainfluenzavirus influenzae* and observed that both are ranked among the top contributors when comparing delta prevalence and SHAP values (Fig. 4D). Additionally, CagY and Hemagglutinin appeared to represent the leading hits for *Helicobacter* and *Betainfluenzavirus* respectively (Fig. 4E).

## Discussion

In this study we characterized systemic antibody repertoires of 321 patients from the CORSA cohort, including colonoscopy-negative controls, adenoma and CRC, against ∼357,000 different antigens. We aimed to determine whether selective group-specific signatures were present at antigenic, protein, and species levels.

Since CRC has been associated with both local and systemic immune alterations (30,31), we hypothesized that disease-associated changes would be reflected in humoral responses against microbial antigens. We observed a global similarity of antigen repertoires across the three clinical groups, indicating that individual immune history is the main driver compared to between-group differences. Similar effects have been reported in IBD, where no global alterations in antibody repertoire composition have been observed. The potential mechanisms underlying this finding should be discussed here. This observation may reflect the fact that the phage libraries used here included a high number of antigens derived from common microbes present as commensals in the gut microbiota (23). Therefore, many antibody responses may represent a shared immunological background shaped by previous exposure to commensals, widely distributed across the population independently of disease-specific responses. This interpretation is further supported by the homogeneity of the cohort, with samples collected within the same geographical area and similar dietary patterns.

In addition, the phage library includes antigens from widely prevalent infectious agents, such as Epstein-Barr and Cytomegalovirus viral antigens (32,33). These were shown to produce long lasting immune responses in a large fraction of the adult population, which may contribute to certain extent to the observed stability of the antibody repertoires. Overall, these findings suggest that a clinical condition such as adenoma and CRC do not result in global remodeling of antibody responses against microbiota but rather selective alterations of immune reactivity against specific microbial antigens, as observed in IBD and PSC (24,25). This is consistent with previous metagenomic studies showing that overall gut microbiota diversity is largely preserved across disease groups, whereas the primary differences are driven by shifts in the relative abundance of specific taxa rather than by their presence or absence. (34).

Despite the global stability in the present study, prevalence-based analyses showed that limited number of antigens were present at different prevalence between controls, adenoma and CRC. This might imply that disease-related humoral alterations are limited to few antigens rather than reflecting broad antibody repertoire-wide shifts.

Interestingly, many human viruses showed at least one antigen to be among the top hits when comparing prevalences across the three groups. For example, *Betainfluenzavirus influenzae* (seasonal Influenza B virus) Hemagglutinin localized antigens were more prevalent in controls and adenoma compared to cancer, *Rhinovirus B* polyprotein was targeted more in controls compared to adenoma, and Herpesviruses proteins (EBV, *Cytomegalovirus*) were differentially detected in the three groups: *Cytomegalovirus* high-prevalence antigens, polyprotein, and tegument protein were detected more prevalently in cancer, while EBV nuclear antigens were detected more often in controls and adenomas. Overall, these signals may reflect general immune background within the cohort. While they do not provide evidence for a strong association with either disease or non-disease status, they indicate trends that warrant further investigation.

For this reason, we aggregated the antigenic signals to lower resolution levels by mapping the peptides to the species and proteins. This provided a better interpretation of biological patterns compared to single antigen analyses. Notably, after statistically testing the aggregated data, few species and protein specific candidates emerged to be relevant for either one or the other group.

One of the most relevant hits was *Helicobacter pylori*, which antigens were consistently detected at higher prevalence in CRC compared to controls and adenoma. *H. pylori* is a pathogen known for its role in causing chronic gastric infection and promoting the development of gastric cancer and its association with CRC (35). Most of the detected signals of varying prevalence was due to two CagPAI genes, namely CagA and CagY. CagPAI is a pathogenicity island including 31 open reading frames (ORF) which encode all the structural and functional elements needed to assemble the Type IV Secretion System (T4SS). This is a machinery that *H. pylori* use to inject the effector protein CagA in the cells of the gastric and intestinal epithelium (36). CagA is an established oncogene which was observed to directly induce DNA double strand breaks (DSB), rearranging cytoskeleton elements and disrupting cell-to-cell tight junctions (37). CagY is one of the biggest proteins encoded in CagPAI, as it is part of the membrane-spanning core of T4SS, and was observed to be an important immunomodulatory element able to tune host inflammatory response (38). The higher prevalence of these components in CRC could indeed point towards an association between *H. pylori* CagPAI-positive strains and higher immune activation in patients with cancer compared to non-cancer, in line with previous observations (39). In addition, the adenoma group showed an intermediate pattern, with higher Cag antigen prevalence compared to controls but lower to CRC. This suggests a potential trend of CagPAI-directed immune reactivity across cancer disease progression.

Contrarily, we found that Influenza B virus (*Betainfluenzavirus influenzae*) was the main hit for both controls and adenoma when compared to CRC. Influenza B virus is one of the main seasonal flu viruses which are also associated with severe respiratory diseases such as bronchitis and pneumonia (40). The antigens we detected to be more prevalent in controls and adenoma were mostly mapped on the Haemagglutinin protein. Hemagglutinin is a surface glycoprotein involved in recognition and binding of host-cell receptors, mediating viral entry in the host organism. It represents one of the most immunogenic targets and therefore it is used to produce seasonal flu vaccines (41). Notably, we found that the antibodies directed against Hemagglutinin antigens in our experiment were mostly detected in conserved domains of the protein, which may explain the high level of cross-reactivity to other Influenza B strains. The higher prevalence of hemagglutinin-directed antibodies in controls and adenoma compared to CRC has to be considered carefully, as a possible difference in the target population of immunological background and vaccination history. Interestingly, a previous epidemiological study has reported associations between influenza vaccination and reduced CRC risk in specific patient population, suggesting a higher relevance of influenza vaccine with broader immunomodulatory effects impacting also on other diseases (42). However, associations observed in this study remain observational and do not provide experimental evidence for a direct role of Hemagglutinin and Influenza B virus in CRC protection.

Machine learning prediction using antigenic presence and absence data supported the already mentioned global stability of antibody repertoires, showing no appreciable classification performance. Nevertheless, a slightly higher performance of the model predicting cancer versus controls or cancer versus adenoma and controls combined pointed to a possible presence of few signatures which could separate cancer from non-cancer patients.

### Strengths and limitations

Leveraging a balanced design of a well-characterized geographically homogeneous cohort (including 321 samples which were sex-age matched across three distinguished clinical groups including colonoscopy-negative controls, adenoma and CRC) allowed systematic comparison of antibody reactivity across the cancer-development axis, while minimizing confounding factors. Furthermore, we made use of a wide and comprehensive PhIP-Seq library, including more than 357,000 antigens belonging to nearly a thousand species of bacteria, viruses, and animal/plant allergens. This large antigenic space allowed us to investigate antibody repertoires of our cohort at an unprecedented scale compared to classical immunological approaches as ELISAs or peptide arrays. Additionally, the analytical framework used here enabled not only to test singular antigens associations, but also protein and species level aggregation for a more biologically consistent interpretation. Thus, we were able to obtain both a global overview of antibody reactivity within clinical groups and specific trends at single antigen, protein and species levels.

Regarding limitations, the lack of strong associations between antigens and groups was partially due to the design of the library itself. As opposed to untargeted/exploratory methods such as metagenomics, PhIP-Seq can only detect what has been included *a priori* in the phage library. The library used here included mostly microbial commensals derived antigens from population scale metagenomics (23,43), while there was no specific enrichment performed for cancer associated pathogens/virulence factors. Therefore, potential disease-associated signals may have been diluted by the background of stable, population-wide microbial reactivity. Nevertheless, PhIP-Seq remains the most powerful method enabling simultaneous screening for wide antibody responses against a high number of antigens, including a heterogeneous number of proteins derived from commensal microbiota, viruses, toxins and allergens.

Two additional limitations are inherent to the PhIP-Seq data itself. First, PhIP-Seq it is not a quantitative method but can only indicate whether an antigen is enriched or non-enriched in the blood of patients. This means that we cannot determine the difference in relative abundance of specific antigens between the groups, as is commonly done with metagenomics and 16S sequencing data. However, the binary data we worked with could still highlight interesting patterns across samples and groups. Second, our PhIP-Seq implementation detects epitopes that span from 54 to 64aa, which primarily represent linear epitopes and smaller secondary structures (23), but are insufficient to capture conformational epitopes that arise through natural protein folding of amino acids more than 54/64 amino acids apart. The ratio of linear to conformational epitopes is experimentally difficult to determine, with estimates as low as 10% for linear epitopes (and conversely 90% for conformational epitopes) (44). Ultimately, given the scale of the library, comprising approximately 357,000 antigens, even if only a small portion of the total humoral responses were captured, this would still represent a broader antigenic space than shown with any other known immunological methods, such as ELISA and peptide arrays.

### Outlook

Future efforts may focus on increasing the sample size, which would increase the statistical power for detection of antigenic associations. The inclusion of an independent external cohort would also be essential to validate our findings, allowing for instance to investigate the impact of different geographical backgrounds on antibody repertoires. In addition, the development of a new cancer-focused PhIP-Seq library enriched in Bacteria and Viruses proteins which have been previously associated with cancer may improve the detection of specific signatures of humoral responses of CRC/Adenoma patients. Finally, integrating PhIP-Seq data with other Omics approaches, such as metabolomics or metagenomics, may add different layers of information and provide more solid biological insights and interpretation.

## Methods

### Study design

#### CORSA study

CORSA is an ongoing multicenter study recruiting participants in cooperation with the province-wide CRC screening program “Burgenland PREvention trial of colorectal Disease with ImmunologiCal Testing” (B-PREDICT) using a fecal immunochemical test (FIT) as an initial screening (26,45). FIT-positive tested participants received a complete colonoscopy and were invited to take part in CORSA. Additional participants were recruited at four hospitals in Vienna. CORSA includes men and women aged between 30 and 90, and excludes patients diagnosed with hereditary CRC syndromes or with any previous cancer history. Plasma samples, questionnaires and written informed consent were obtained from all study participants. Information on demographic (e.g. age at diagnosis, weight, height) and lifestyle factors (including diabetes status, alcohol consumption, and smoking status) was obtained through self-assessment using the basic CORSA questionnaire. All CRC patients were diagnosed as histologically confirmed, sporadic CRC, stage I-IV. Patients diagnosed with adenomatous tubular polyps larger than 1 cm, adenomatous tubulovillous polyps, and/or adenomatous villous polyps were included in the adenoma group. Control patients were population-based, FIT-positive, and colonoscopy-confirmed to be free of polyps and/or CRC. EDTA blood samples were obtained at time of colonoscopy, prior to surgery or any radio- or chemotherapy, processed within 4 h following standardized protocols and stored at - 80°C until further processing. Samples used in this study were then selected from the biobank and matched by age and sex across the three clinical groups (controls, adenoma and CRC).

#### Assessment of clinical data and follow-up

Clinical data were extracted from medical records and processed in a structured database following standardized documentation guidelines and according to the General Data Protection Regulation (GDPR). The CORSA database comprises structured information on diagnosis, treatment, histology, progression (recurrence and metastasis), and survival data. Follow-up on clinical data is regularly performed.

#### PhIP-Seq library composition

In this study, three already published antigen libraries comprising a total of 357,192 distinct oligopeptides variants were pulled together: a 244,000-variant library (“Agilent”), a 100,000-variant library (“Twist”) and a 13,192-variant coronavirus library (“Corona2”) (23,27,28). The first library included 244,000 antigens covering 28,668 proteins, mainly derived from microbial strains. In particular, the library covered proteins derived from commensals strains belonging to human microbiota, probiotics, antibody-coated bacteria and gut pathogens. Additional proteins included in the library were retrieved from public databases such as the Virulence Factor Database (VFDB) and Immune Epitope Database (IEDB) (46,47). The second library included 100,000 antigens belonging mainly to allergens, bacteriophages and bacterial strains. For this, six databases were used for protein sequences collection, namely the Immune Epitope Database (IEDB), WHO/IUIS Allergen Nomenclature Database (48), AllergenOnline (49), Allergome (50), AllFam (51) and the Structural Database of Allergenic Proteins (52). Third and last library was the coronaviruses library, which included all seven known human coronaviruses and 49 additional animal coronaviruses.

### Immunoprecipitation and Next Generation Sequencing

PCR plates used for bead transfer and washing were blocked overnight at 4°C with bovine serum albumin (BSA; 30 g/L in Dulbecco’s phosphate-buffered saline). For IP, diluted phage/antibody mixtures were supplemented with BSA (2 g/L), and the phage wash buffer contained 0.1% IPEGAL CA-630 (Sigma-Aldrich, catalog no. 13021). Serum IgG (3 μg, quantified by ELISA) was incubated with phage libraries at a 4,000-fold coverage per library variant, following optimization experiments to determine optimal antibody and phage input amounts. The 244,000-variant library was combined with a 100,000-variant 200-nt library at a 2:1 ratio. Phage/antibody mixtures were incubated overnight at 4°C on an overhead rotator in 96-well plates. Protein A/G magnetic beads (Thermo Fisher Scientific, catalog nos. 10008D and 10009D) were prepared according to the manufacturer’s instructions, and added to samples (40 μL, 1:1 ratio). After a 4-hour incubation, beads were transferred to PCR plates and washed twice using a Tecan Fluent 1080 liquid-handling robot equipped with filter tips. Pooled Illumina amplicon sequencing libraries were generated using Q5 polymerase (New England Biolabs, catalog no. M0493L). PCR amplification consisted of three sequential reactions: PCR1 using library-specific primers, PCR2 using 10-nucleotide unique dual index primers (purchased from IDT), and PCR3 using pool amplification primers. PCR3 products were gel-extracted and purified twice using QIAquick gel extraction and PCR purification kits (Qiagen, catalog nos. 28706 and 28106). Sequencing was performed with custom R1/R2 primers (44/31 nt) on an Illumina NovaSeq 6000 platform using paired-end sequencing.

### Raw data preprocessing, normalization and antigens enrichment computing

Raw fastq files were analyzed through a standardized pipeline assessing the number of enriched oligopeptides per sample – the reactive antigens. Reads were aligned against the PhIP-Seq oligopeptides sequences using Bowtie2 (53), in single-end mode and using *--end-to-end* setting, keeping only the best alignment. Per-peptide coverage was calculated using SAMtools (54), by filtering the alignments according to their edit distance, previously obtained through Bowtie2. Final oligopeptide count was derived as the maximum observed coverage depth across each peptide sequence and saved in tabular format. Oligopeptides counts were then normalized with a similar approach used previously (55). Specifically, the normalization approach consists in comparing, for each sample, the peptide counts to the counts obtained by the input (the sequenced library before immunoprecipitation with the antibodies): These counts are used to estimate the null distribution of the counts expected in absence of antibodies. We fitted the null distribution using a Zero-Inflated Generalized Poisson (ZIGP) model. Oligopeptides enrichment was then assessed by comparing the observed output counts against the fitted peptide counts distribution, and the p-value value was calculated as the right-tail probabilities under the fitted ZIGP null model. P-values were adjusted for multiple hypothesis testing using Holm method and peptides which did not reach enrichment were discarded. Fold-change from the input counts were calculated per each peptide as the ratio between each peptide counts per million (cpm) the corresponding value in the input. For each sample, peptide-level presence/absence profiles, enriched peptides counts and foldchanges were aggregated across all samples to construct three oligopeptide-by-sample matrices: a binary presence/absence matrix, a corresponding count matrix and fold-change matrix. Prior analyzing the data, we removed 10 samples which accounted for less than 30 enriched oligopeptides.

### Data analysis

#### Assessing Sex/Age matching

Age and sex were previously observed to influence PhIP-Seq data analysis, since antibody repertoires can shift according to both age and sex differences. Matching across all group pairwise comparisons was assessed by running a logistic regression with group status as the outcome and age and sex as predictors: p-values were obtained by likelihood-ratio test by comparing the full model with an intercept-only model used as null model.

#### Peptide prevalence, relative abundance and reactivity breadth

For each group we computed the peptide prevalence and a per-protein breadth reactivity score. The peptide prevalence was simply assessed dividing the occurrences (n) of the same peptide antigen in each group divided by the total number of samples (N) in the given group.

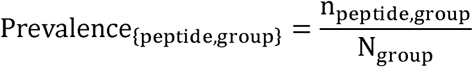

We also defined the reactivity breadth by obtaining for every protein in the library the fraction of total peptides/antigens that were bound in each sample. The reactivity breadth of a protein is higher the more antigenic fraction of that protein is detected by antibodies in each sample:

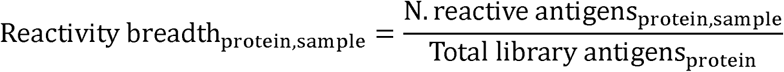

#### Groups within-sample and between-sample diversity

Using the peptide presence/absence and count profiles, we performed diversity analyses to evaluate whether antigen profiles differed across diagnostic. Alpha diversity was assessed from peptide count profiles using Shannon diversity index, measuring the within-sample diversity in antigen reactivity. Differences in the alpha diversity across groups was then assessed by Welch’s t-test. To assess the between-sample diversity and observe if this deviates across groups, we performed beta diversity analysis using Jaccard dissimilarity index on presence-absence sample matrix to calculate sample distances. We visualized the resulting matrix by through Principal Coordinate Analysis (PCoA) and running PERMANOVA to assess group-level differences.

We also applied beta-diversity after aggregating peptides to the species taxonomic level for each sample, generating a “species” presence-absence matrix. The presence or absence of a species was assessed per sample when at least 1 peptide belonging to that species was enriched. We assessed in the same way if group-level differences were present by running PERMANOVA.

#### Taxonomic-level prevalence aggregation: the delta framework

An interesting question in the analysis of high-dimensional data with hierarchical structure is how to test statistical hypotheses at different levels of that structure. PhIP-seq data is characterized by an exact taxonomic structure, as well as by multiple variables that group individual peptides into higher-order sets (such as allergens or flagellins), which we refer to as annotations. To address this question, we developed a statistical analysis framework called DELTA (Differential Enrichment via Label-permutation Testing of Annotations). DELTA computes per-peptide Z-scores using the signed root likelihood-ratio test statistic, aggregates them using the maxmean approach of Efron and Tibshirani (56), and assesses statistical significance through subject-level label permutations, which preserve the complex dependence structure among peptides.

Because this analysis is primarily exploratory, we did not apply false discovery rate (FDR) correction. Instead, we identified the relevant biological signal in each comparison using three parameters, filtering hits based on:

- effect size, represented by the normalized aggregated maxmean statistic;
- the number of peptides belonging to a given annotation; and
- the maximal prevalence of peptides within an annotation.

Specifically, for CRC versus controls and CRC versus adenoma, we retained hits with at least 30 peptides per annotation, a |standardized maxmean statistic| ≥ 1.5, and a maximal prevalence ≥ 0.1. For adenoma versus controls, we adjusted the effect-size threshold (|maxmean| ≥ 1.8) to account for the greater variation in the coefficient distribution.

#### Machine learning predictions

We leveraged machine learning for predicting the group category by using the antigens presence/absence data as features. We trained and optimized random forest models to predict CRC versus adenoma, adenoma versus controls and CRC versus controls. We used a Nested Cross-validation scheme slitting the data in training set (80% of the data) and test set (20%) using 5 inner and 5 outer folds: The inner folds were used for training the model, feature selection and hyperparameters tuning. Features were selected based on how much were they informative, using a lasso regression within each inner fold. We then trained a random forest classifier, optimizing the hyperparameters with Bayesian search and evaluating the model. Outer folds were then used to test the model, using a random forest classifier with the optimized hyperparameters selected within the inner folds. We then averaged the results for all outer folds and computed the areas under the curve (AUC). A second model was trained in the same way to recognize CRC versus adenoma and controls and then controls versus CRC and adenoma with the same setup.

#### Multi-sequence alignment of Hemagglutinin vaccine variants

We downloaded five Influenza B virus Hemagglutinin variants that were widely used in Europe between 2002 and 2015 as vaccine (“ABL77178 B/Hong Kong/330/2001 2001/04/09 HA”, “ABO37784 B/Ohio/1/2005 2005/02/23 HA”, “AFH57953 B/Wisconsin/01/2010 2010/02/20 HA”, “AGL06036 B/Massachusetts/02/2012 2012/03/13 HA”, “CAH04474 B/Shanghai/361/2002 2002// HA”. We performed a multi-sequence alignment using mafft v7.526 (2024/Apr/26) (57). Alignments were plotted using python3.

### Ethical aspects

Written consent was obtained from all study participants, and all studies were approved by the corresponding Institutional Review Board. Compliance with the 1964 Declaration of Helsinki, the Austrian Drug Law (Arzneimittelgesetz, AMG) and the requirements of Good Clinical Practice of the European Community (CPMP/ICH/135/95) will be ensured. The CORSA study was approved by the institutional review boards (EK 33/2010 and EK 1160/2016).

### Data availability

Raw sequencing data is available at https://doi.org/10.7910/DVN/CFETZP.

### Code availability

Data analysis was performed through in-house scripts in python and R. Code is available at https://doi.org/10.5281/zenodo.23364597. Packages performing Delta statistics can be found at https://github.com/Vogl-Lab-Research/phiper and machine learning method used for performing predictions at https://github.com/csReynaB/phipml.

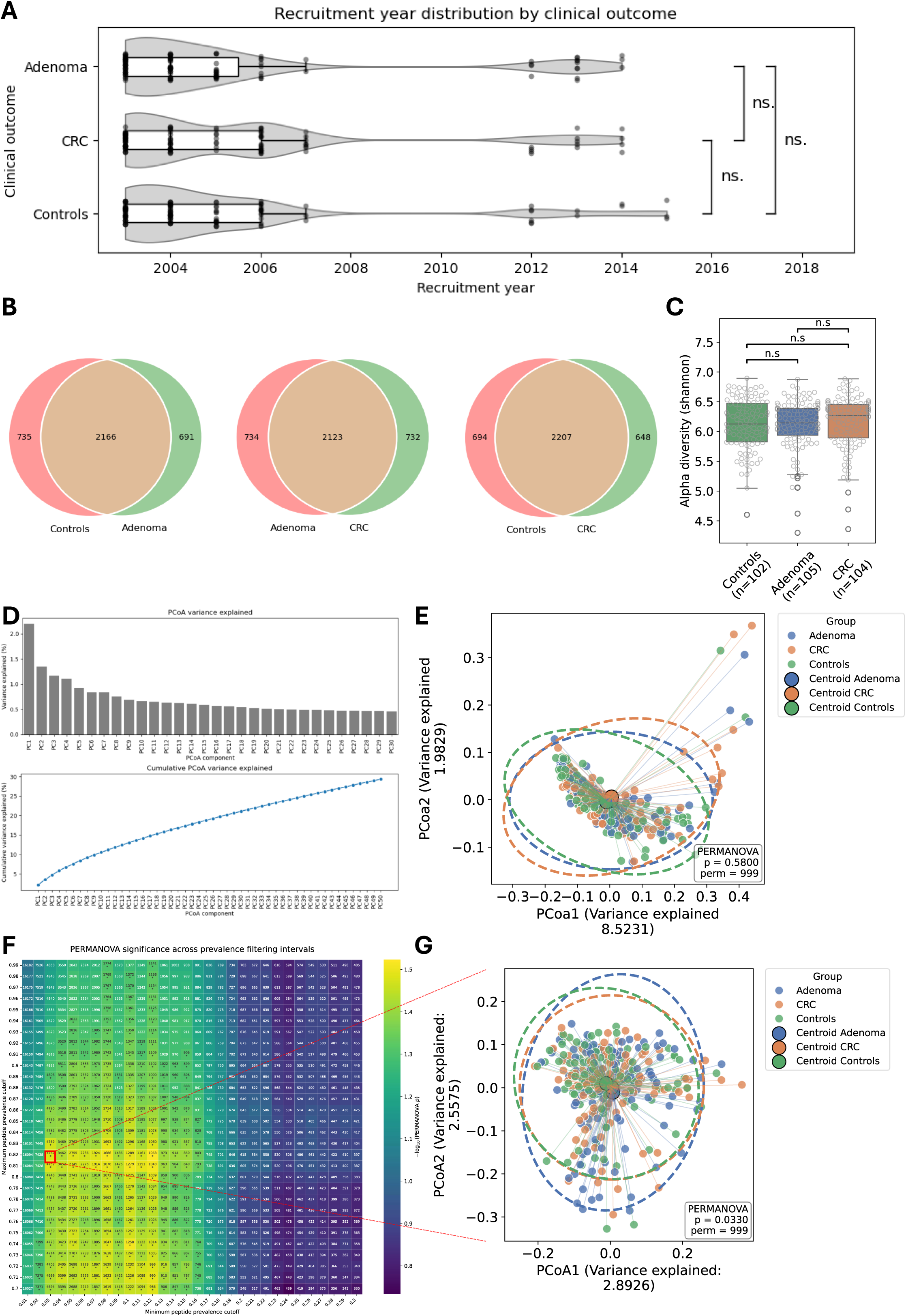

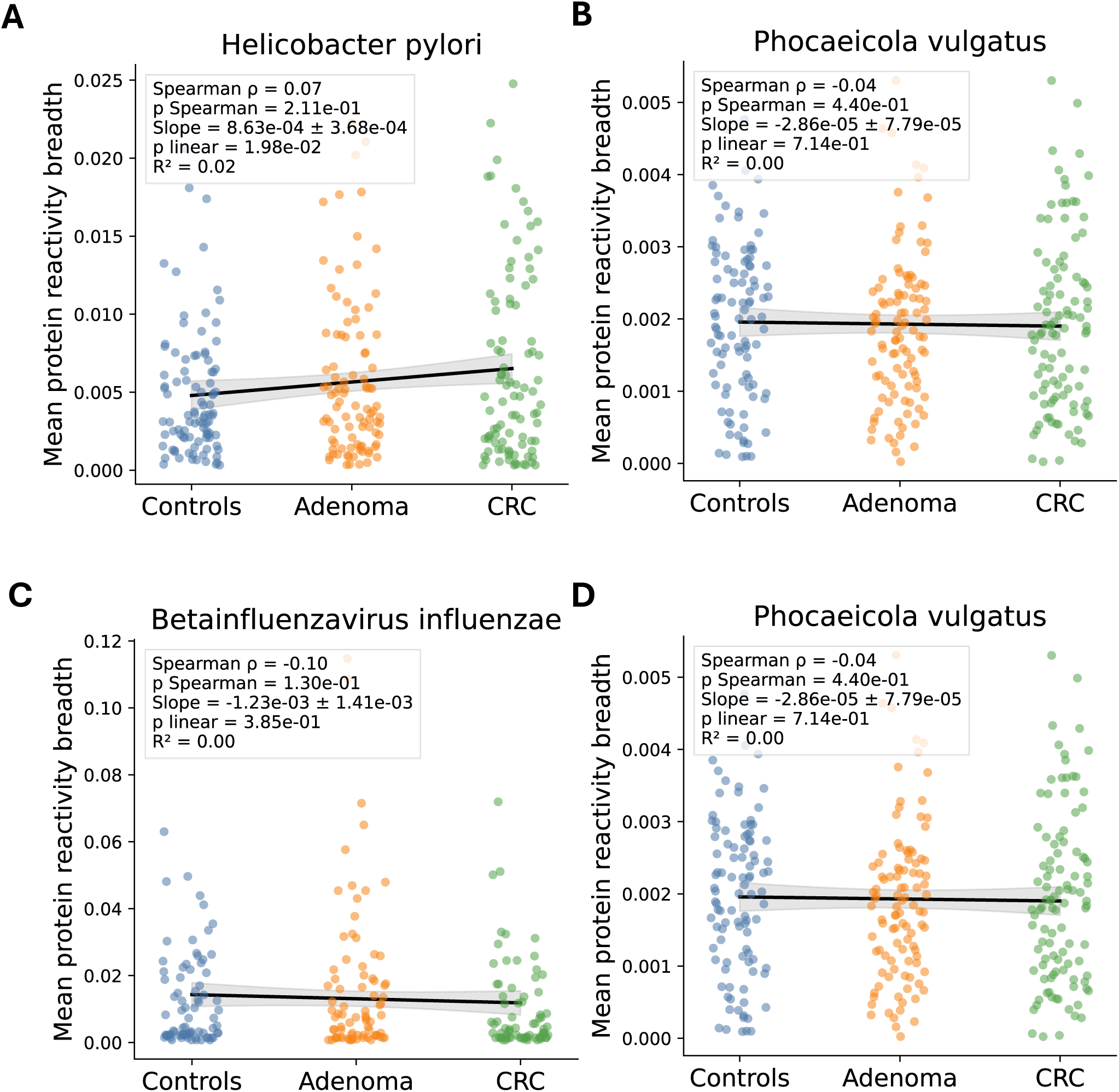

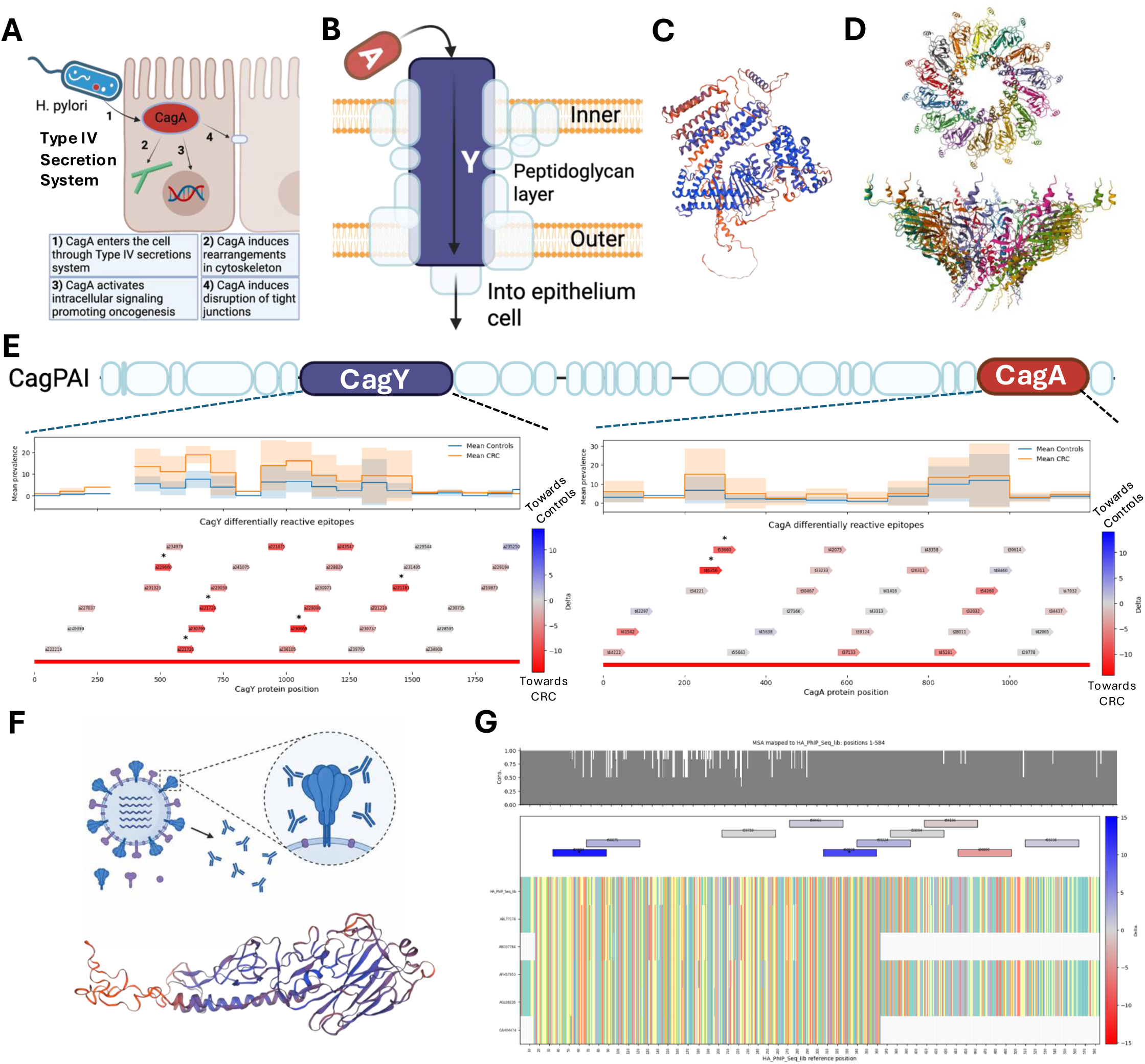

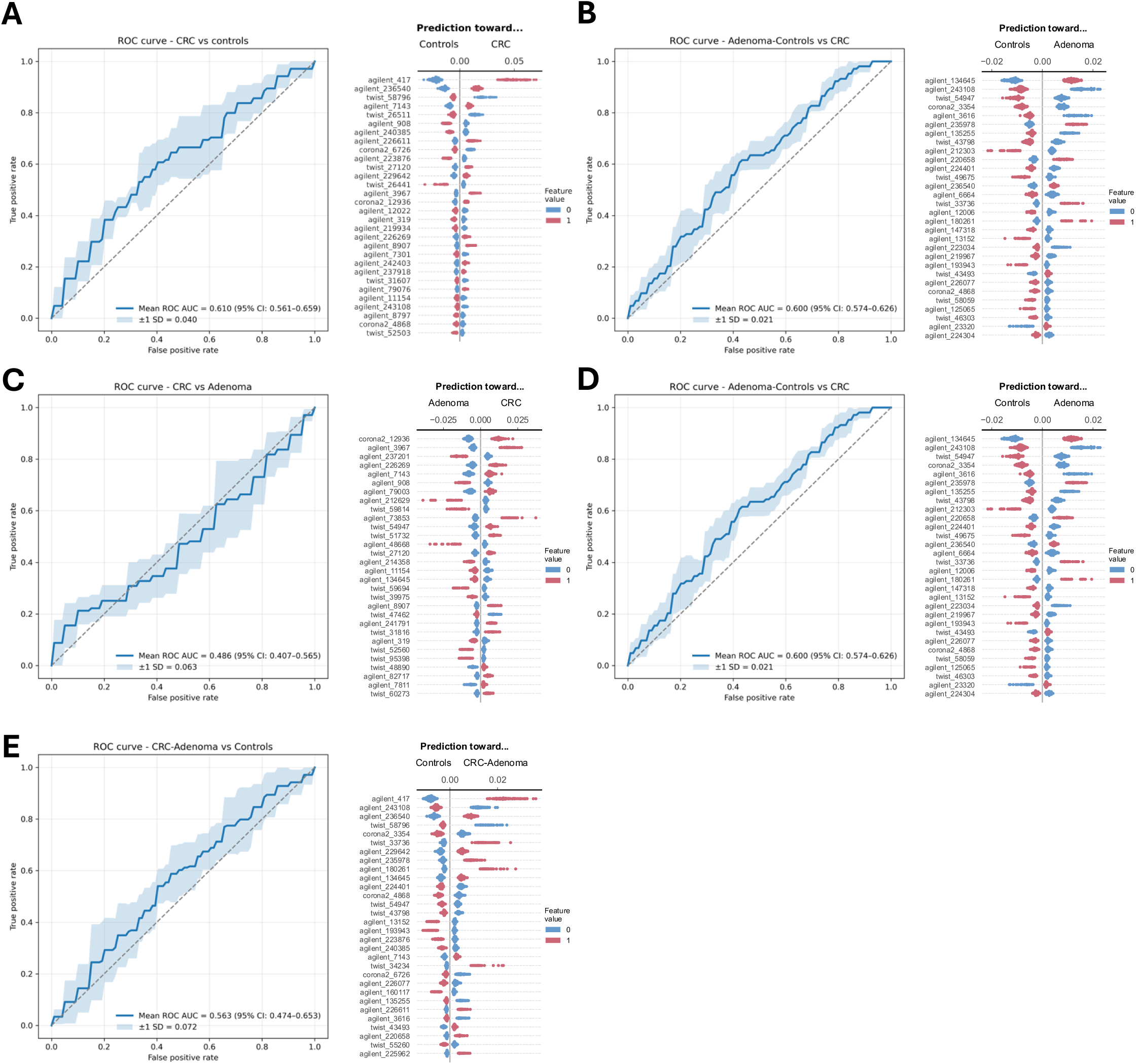

## Supporting information

Captions and text for supplementary figures and tables

## Funding declaration

The project was funded by European Research Council [ERC] Start Grant EarlyMicroAbs [project number 101075733] and Horizon Health consortium “ID-DarkMatter-NCD” [PN: 101136582].

## Authors contribution

G.I., S.B., A.G. and T.V. conceived and designed the study. S.B., A.G., J.P. and A.M. recruited the study participants and collected the sample metadata. M.P. performed sample processing and PhIP-Seq experiments. G.I. and C.R.B. performed the bioinformatic and statistical analyses, with statistical support from M.F.K. I.B. contributed to data interpretation and study design discussions. G.I., S.B. and T.V. wrote the manuscript. T.V. and A.G. coordinated and supervised the study. All authors revised the manuscript and approved the final version.

## Acknowledgements

T.V. gratefully acknowledges support from the European Union (European Research Council [ERC] and Horizon Health consortium), as well as support from the EMBO YIP program (#6165).

We kindly thank all individuals who agreed to participate in CORSA. Furthermore, we thank all cooperating physicians and students.

## Notes

https://doi.org/10.7910/DVN/CFETZP

https://doi.org/10.5281/zenodo.23364596

### Competing Interest Statement

The authors have declared no competing interest.

