## Supplementary material for "Systemic antibody repertoires target specific microbial antigens in patients with colorectal cancer": Captions and text for supplementary figures and tables

### Supplementary files

#### Supplementary Table 1

Supplementary Table 1: Samples metadata including age, sex and clinical group.

#### Supplementary Figure 1

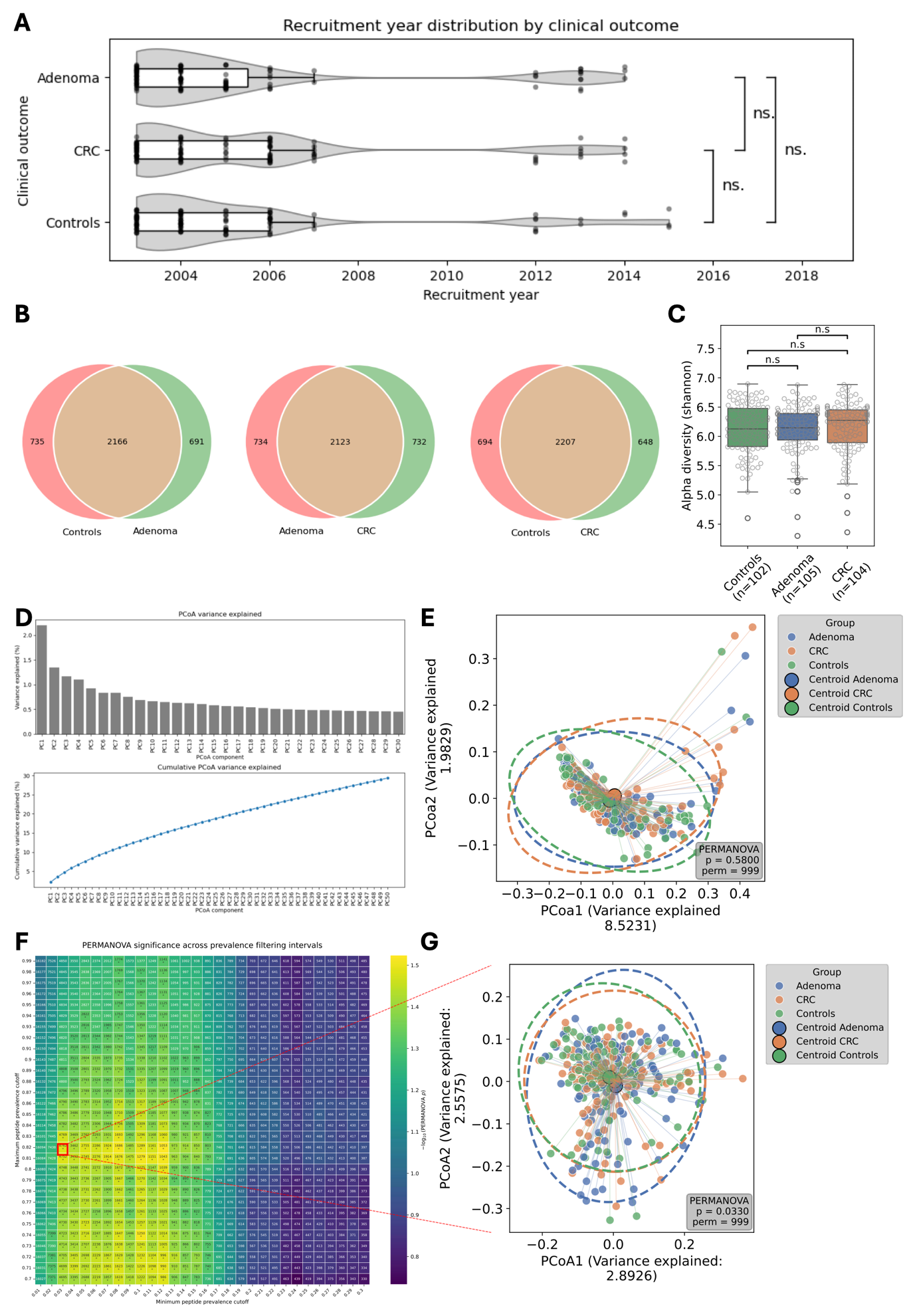

Supplementary Figure 1: Global stability of antibody repertoires remains even if we select antigens at 5-100% prevalence interval, if we aggregate them to species, or if we compare CRC versus non-CRC group (Adenoma + Controls).

A. Comparison of recruitment year distribution within the different clinical groups.

B. Venn plot showing core and accessory antibody responses in each comparison with antigens present in 5-100% prevalence interval across the three groups.

C. Alpha diversity boxplot using Shannon index on the three groups.

D. Bar plot showing the variance explained for each dimension in the Principal Coordinate Analysis (PCoA1 and PCoA2 shown in Figure 1F) absolute (upper plot) and cumulative (lower plot).

E. PCoA using antigens presence-absence data and testing for differences within the cancer versus non-cancer comparison.

F. Heatmap reporting PERMANOVA test results on all possible subsets of peptides obtained by applying all combinations of prevalence-based filtering thresholds, with lower cutoffs ranging from 0 to 0.30 (0-30% lower cutoff) and 0.7-0.99 (70-99% upper cutoff). Each result is reported in the heatmap encoding the PERMANOVA p-value as color, number of peptides used post filtering and whether the test showed to be significant (no FDR used for this exploratory analysis).

G. Selected subset showing PERMANOVA pvalue <= 0.05 with a number of 4,752 antigens was used for showing beta diversity (PCoA1 versus PCoA2).

#### Supplementary Table 2

Supplementary Table 2: Table showing difference in prevalence between antigens in each group pair comparison, tested with Fisher test and corrected for multiple hypothesis testing (only significant hits prior correction shown).

#### Supplementary Table 3

Supplementary Table 3: Summary table showing the results statistical analysis of antigenic prevalence aggregation to species, genus and protein annotation level (delta framework, Methods).

#### Supplementary Figure 2

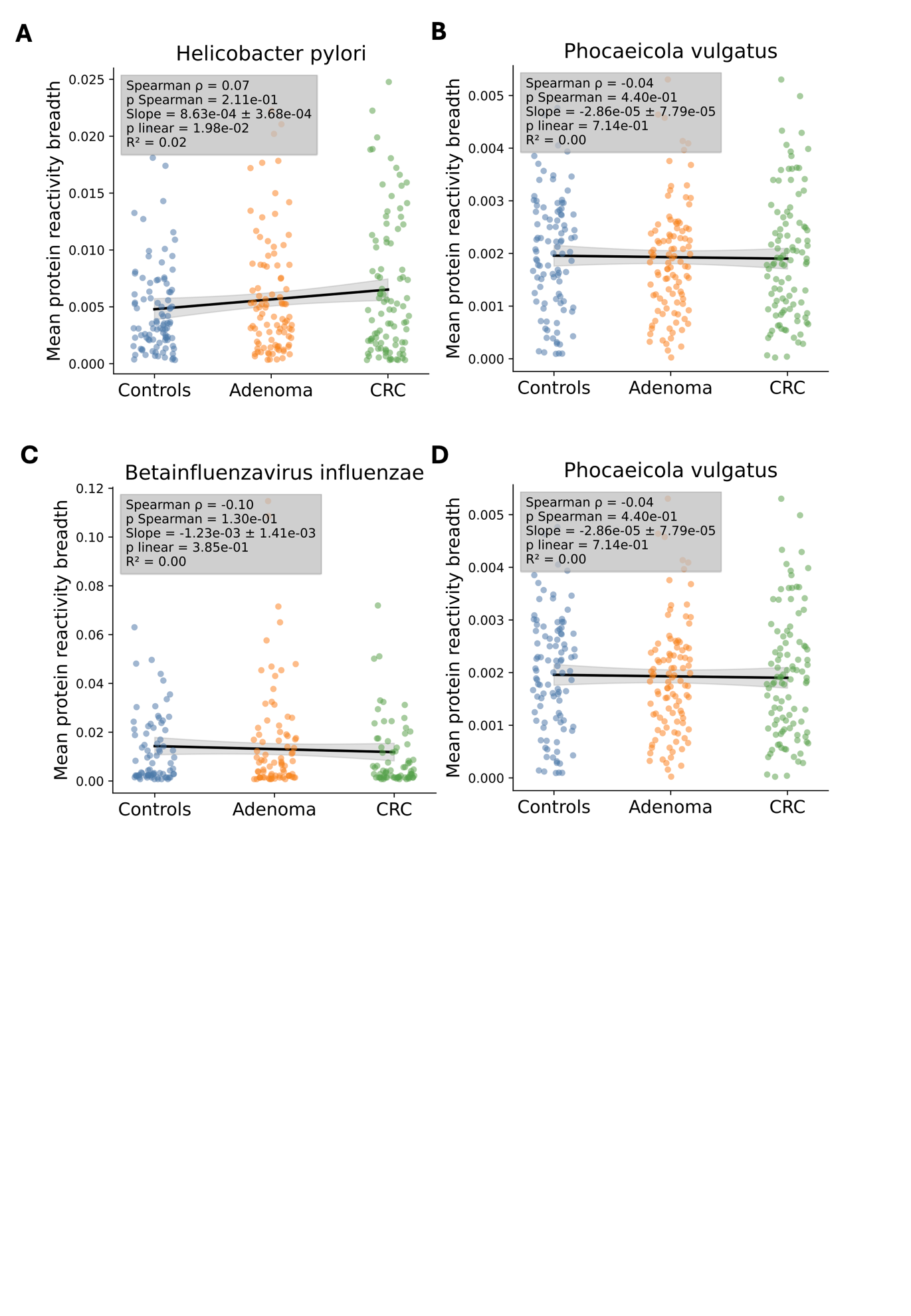

Supplementary Figure 2: Comparing protein mean reactivity breadth for the main microbial hits obtained after delta framework statistical analysis for each group showed an increasing trend only for Helicobacter pylori.

A. Comparison of protein mean breadth between clinical groups for Helicobacter pylori.

B. Comparison of protein mean breadth between groups for Subdoligranulum sp. APC924/74.

C. Comparison of protein mean breadth between groups for Betainfluenzavirus influenzae (Influenza B virus).

D. Comparison of protein mean breadth between groups for Phocaeicola vulgatus.

#### Supplementary Figure 3

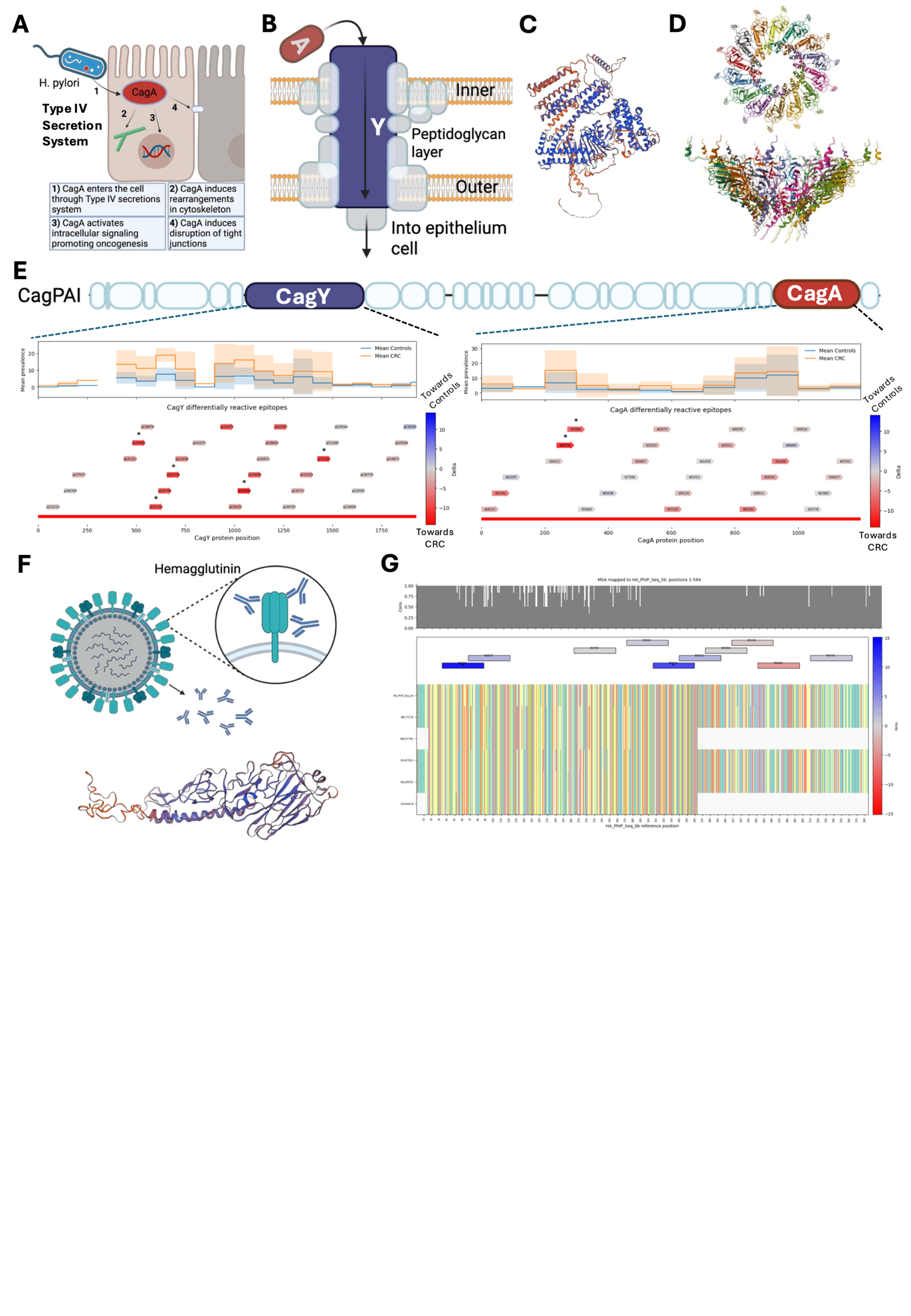

Supplementary Figure 3: The Cag pathogenicity island and Hemagglutinin as potential differently prevalent antigens CRC and controls.

A. Schematic overview of the mechanism of action of Helicobacter pylori Type IV secretion system, observed mainly in gastric cancer.

B. Schematic structure of CagPAI secretion system, highlighting CagY and CagA that were found to be differentially prevalent in CRC versus controls.

C. The structure of CagA based on Swiss prot predictions.

D. The predicted 3D structure of CagY transmembrane element based on UniProt predictions.

E. Schematic view of genomic Cag pathogenicity island (CagPAI) with the antigen’s coverage showed below for CagY and CagA. The color encodes the difference in prevalence (Delta prevalence) between CRC and controls.

F. Hemagglutinin protein from Influenza B virus schematic overview and the structure based on SwissProt predictions.

G. Multi-sequence-alignment of Hemagglutinin proteins from vaccine strains used between 2003-2007 and 2013-2016. The figure highlights the antigenic sequences that were differentially bound in CRC (red) and controls (blue), showing that the most conserved area was the one exhibiting the higher difference in binding prevalence.

#### Supplementary Figure 4

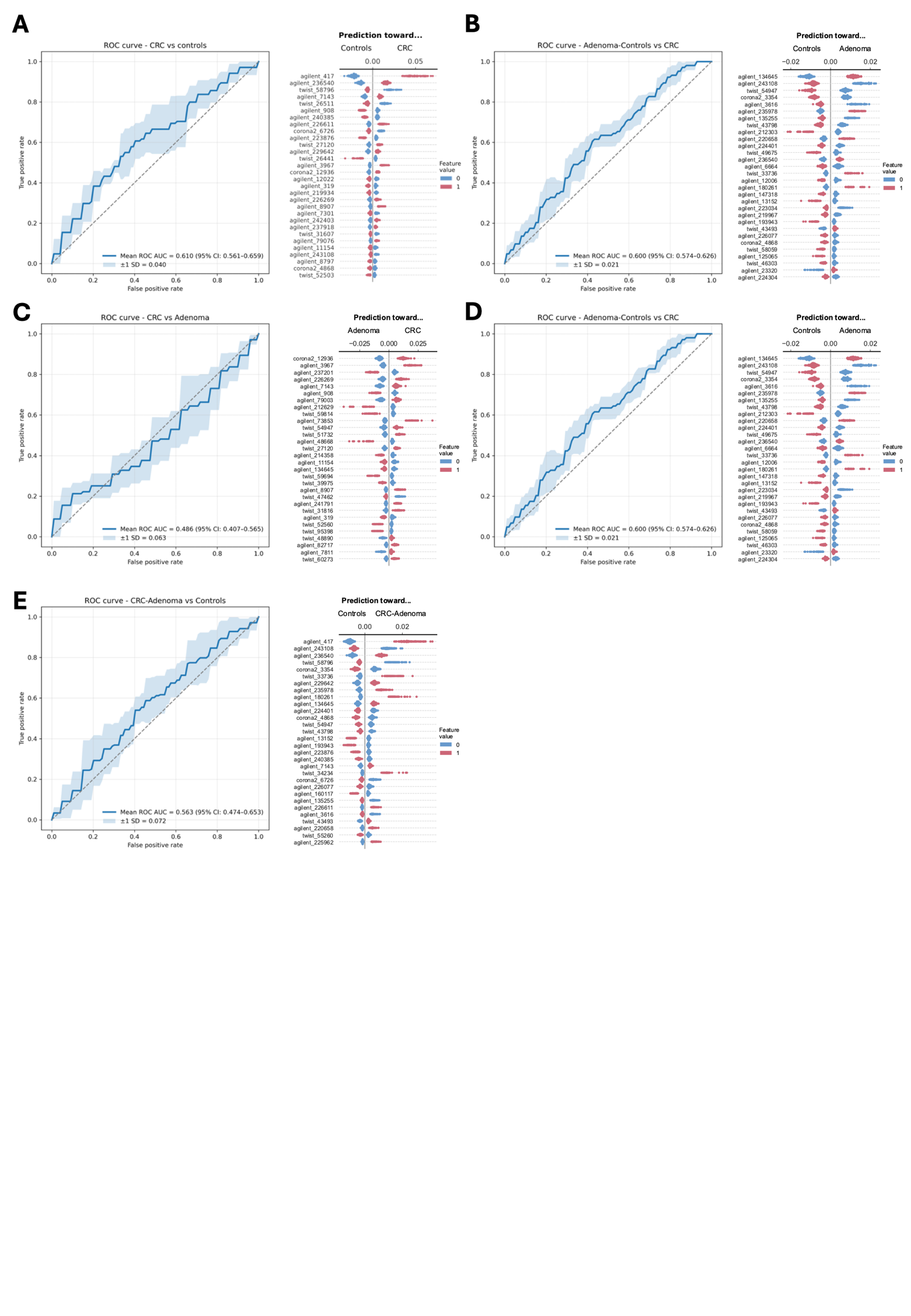

Supplementary Figure 4: Random Forest predictions on each group pair comparison shows no high performance, with a better result when comparing CRC versus Controls or CRC versus non-cancer.

A. CRC versus controls.

B. CRC versus Adenoma-Controls.

C. CRC versus Adenoma.

D. Adenoma versus controls.

E. CRC-Adenoma versus Controls.
